# AI semantics for biomedical data integration

**DOI:** 10.64898/2026.08.03.742514

**Authors:** James McLaughlin, Aleix Puig-Barbe, Arwa Ibrahim, Diego Pava, Zoe May Pendlington, Nicolas Matentzoglu, Elliot Sollis, Amy Foreman, Robert Wilson, Federico Lopez Gomez, Laura Harris, Yeyejide Adeleye, Satwant Kaur, Birgit Meldal, Damian Smedley, Helen Parkinson

## Abstract

Researchers increasingly need to explore hypotheses that span multimodal data across different scales, organisms, and domains. In practice, this requires connecting knowledge across fragmented databases with incompatible APIs and heterogeneous annotation practices. Large language model (LLM) agents can automate this data integration process, but grounding LLM agent outputs in scientifically correct sources of truth remains a significant challenge.

Here we describe our deployment of a novel AI semantics workflow using LLM agents to enable scalable data integration, grounded in biological knowledge in the form of ontologies. Our workflow comprises (1) a multi-agent system curating scientific knowledge across ontologies using the Ontology Lookup Service (OLS) as grounding; (2) an LLM embedding service to enable interoperability between scientific databases by mapping ontology terms; and (3) GrEBI, a knowledge graph and Model Context Protocol (MCP) server enabling LLM agents to conduct cross-cutting, multi-omic biomedical queries.

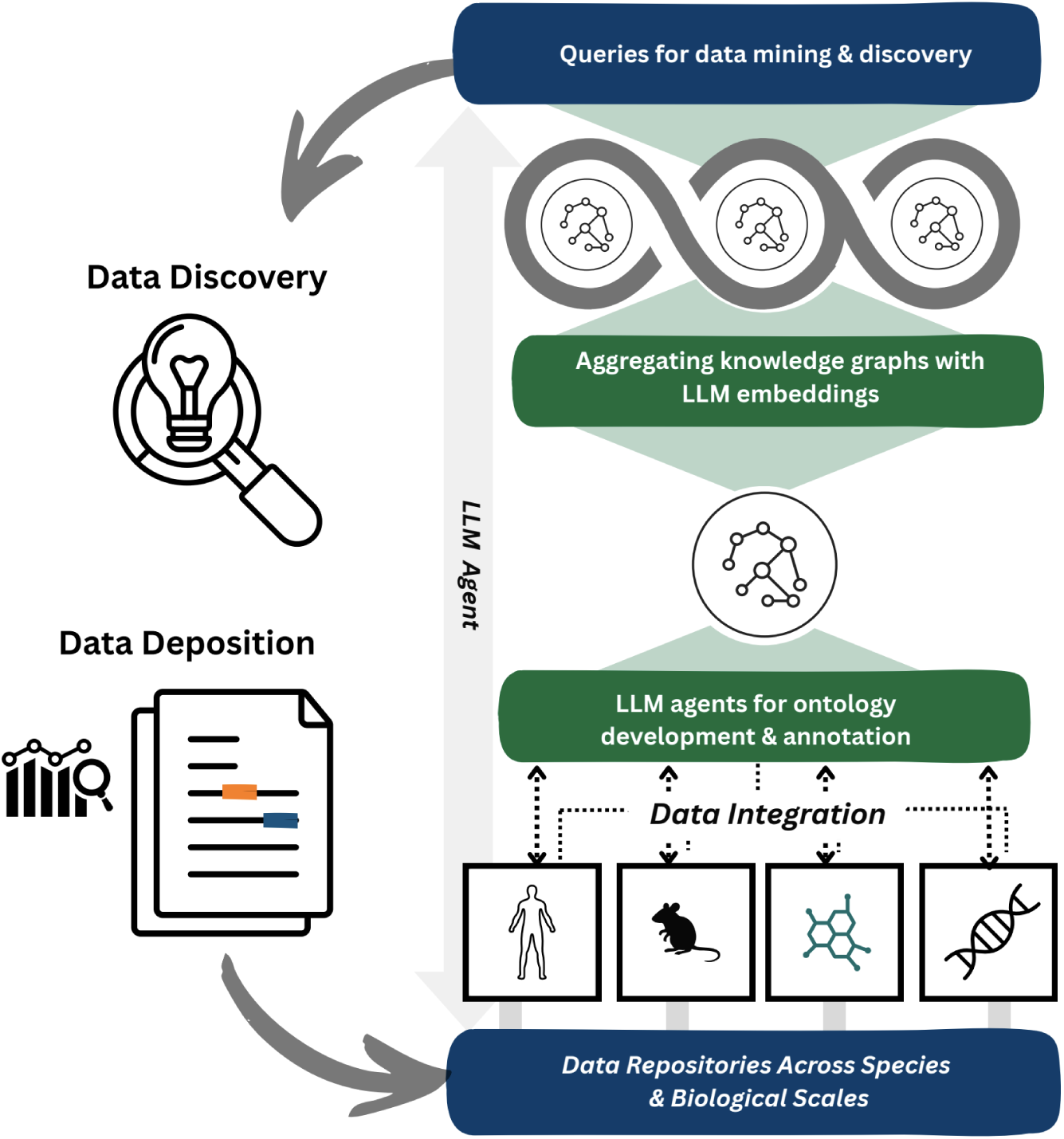

## Introduction

Advances in biomedical research increasingly depend on integrating multimodal data across different scales and organisms to test complex hypotheses. For example, in drug target selection for disease, relevant knowledge can include cohort derived human genetic data, mouse model phenotyping data, and cell-based screening. To enable this integration, efforts such as the Monarch Initiative KG, DisGeNET^1^, Bio2RDF ^2^, ROBOKOP^3^, PrimeKG^4^, and BioCypher^5^ bridge incompatible database APIs and schemas by integrating data into knowledge graphs (KGs). This integration depends on annotation of databases with ontologies, which define shared semantics for biomedical concepts. EMBL-EBI develops the Experimental Factor Ontology (EFO)^6^, an ontology supported by the public-private partnership Open Targets with applications in drug-target prioritisation and validation^7^; Genome Wide Association (GWAS) study trait annotation^8^; transcriptomics^9^; bioassays^10,11^; and bridging to model organisms^12,13^. EFO defines terms for diseases, phenotypes, traits, responses to chemicals and drugs, cells, and anatomy, using standard domain ontologies such as OBA^14^, MONDO^15^ and the Human Phenotype Ontology (HPO)^16^ to enable semantic interoperability across different databases, and local terms where needed.

As scientific use cases increase in complexity, there is a need to scale up semantic methods to enable deeper data integration beyond the scope and granularity provided by existing ontology-based approaches. This need was highlighted by a collaboration between EMBL-EBI, AstraZeneca, Pfizer, and Syngenta evaluating ontology methods to integrate knowledge on rare syndromes with sparse data such as Bronchiectasis and Thrombotic microangiopathy, which identified limitations in current annotation practices. In one example, identifying metabolites or drugs relevant to disease in GWAS Catalog studies was limited by annotation of studies with nonspecific ontology terms e.g. `OBA:2050078 blood metabolite level` and `EFO:0009170 response to tyrosine kinase inhibitor` rather than terms with specific metabolite or drug identifiers such as “ornithine metabolite levels” and “response to nintedanib”, many of which were missing from existing ontologies. In another example, validating human gene-phenotype associations with synergistic mouse models in the International Mouse Phenotyping Consortium (IMPC) database was limited by biologically related annotations not being semantically mapped, such as `EFO:0004309 platelet count` and `MP:0003179 Thrombocytopenia.` In both cases -the need for specific ontology terms, and mappings across biological and semantic distance - manual curation from a domain expert would have been required, which is a limiting factor for scalability.

Large language models (LLM) agents have the potential to address these complex data integration scenarios due to their ability to bridge syntactic and semantic heterogeneity. However, LLMs require grounding to avoid hallucinations ^17^ ^18^. For example, querying the GPT-5 model with *“What are the main phenotypes of Bronchiectasis? Use HPO terms”* returned an unrelated phenotype `HP:0031251 Abnormal subclavian artery morphology` which was incorrectly labelled “Excessive sputum production” in the response (Supplemental S1). This example illustrates that LLMs can bridge semantics and context, but without grounding in knowledge they can return plausible but incorrect results. Despite this challenge of hallucination, LLMs have already been shown to enhance existing ontology-mediated integration methods; for example, Uberon and MONDO employ the Claude-based DRAGON-AI agent^19^ to create new ontology terms needed to annotate data. LLM embeddings can be used with ontologies for Retrieval Augmented Generation (RAG) approaches^20,21^ and to associate previously unconnected concepts across ontologies by embedding similarity^22^. Altogether, there is an opportunity to enable large-scale, accurate data integration with a combined workflow, where the reasoning flexibility of LLMs is complemented by the accuracy and structure of ontologies.

## Results

Here we describe our deployment of a novel AI semantics workflow for biomedical data integration using LLMs and ontologies synergistically, and its application to integrating disease knowledge across species. First, we have redesigned the curation process for relevant datasets, using a multi-agent workflow grounded in scientifically accurate and up-to-date ontology terms via a Model Context Protocol (MCP) server for the Ontology Lookup Service (OLS). We have applied this method to establish new connections from the GWAS Catalog to external databases by using LLMs to curate drug response and metabolite measurement terms at the resolution of specific metabolites and drugs. Secondly, we have scaled up LLM embedding methods for ontology mapping to cover all of the ontologies in OLS, which we have applied to connect semantic and biologically distant concepts from human GWAS studies with mouse models in IMPC. Finally, we have implemented GrEBI (Graphs@EBI), a KG and MCP server to complete the workflow by enabling LLM agents to use the resulting integrated data, demonstrated by exploring similar phenotypes across orthologous human and mouse genes.

### Multi-agent ontology curation with grounding MCP

Curated biomedical databases enable researchers to access structured, standardized data, making information computationally searchable, comparable across studies, and suitable for large-scale analysis. To construct curated databases at EMBL-EBI, publications and submitted data are annotated using standard terms by expert curators; for example, a study referencing “fibrosis” would be mapped to `EFO:0006890 fibrosis` for inclusion in the GWAS Catalog. We have scaled up this manual curation process while minimising the risk of hallucinating ontology terms, by developing an MCP server for OLS to enable LLM agents to access ontologies as a grounding framework to assist biomedical curators. For example, Perea *et al.* (2024)^23^ includes a graphical figure describing the four main pathophysiology parameters of bronchiectasis (infection, inflammation, lung damage, and impaired mucociliary clearance). To map these to ontology terms, a curator entered the following prompt: “Extract bronchiectasis pathophysiology terms from the image. Use the OLS MCP to retrieve ontology IDs and labels.” The agent correctly identified the four key pathophysiology terms from the image and provided a table with mapping to valid ontology identifiers (Supplemental S2). Compared to the previous ungrounded example of requesting the Bronchiectasis phenotypes which returned a hallucinated ontology term `HP:0031251 Abnormal subclavian artery morphology`, in this case the role of the LLM only interprets supplied knowledge, rather than providing any scientific or semantic input. The scientific evidence is provided by the literature, and semantic grounding is provided by the OLS MCP, therefore reducing the risk of hallucination while still benefiting from the flexibility of the LLM to extract knowledge across text and images.

A further challenge in curation is where the required terms for annotation are not available in existing ontologies, due to the need for manual curation by ontology editors. In these cases database curators often substitute missing ontology terms with broader terms which are already available when codifying new or challenging concepts, which limits the granularity for search and data integration. For example, GWAS studies are increasingly making use of high-throughput metabolomics platforms^24^, yet the nonspecific term `OBA:2050078 blood metabolite level` is used to annotate 4494 GWAS Catalog studies (as of August 2026) in the absence of terms for specific metabolites. This lack of specificity in annotation prevents data integration; for example, cholesterol sulfate is involved in cholesterol homeostasis, but as `GCST90615620 Cholesterol sulfate metabolite level` is annotated with the generic `OBA:2050078 blood metabolite level` it cannot be connected to cholesterol metabolism and its related primary lipid metabolism pathways.

To address this issue we have developed a multi-agent workflow grounded using the OLS MCP which enables LLM agents to create missing ontology terms with accurate references to existing ontologies. Our workflow consists of four agents: a co-ordinating agent (main Copilot Coding Agent); an ontologist agent; a curator agent; and an importer agent. The co-ordinating agent reviews the tickets opened by curators and assesses the information provided, generating a plan to address each request. Where the plan requires further evidence or domain validation, the ontologist agent delegates to the curator agent. The curator agent performs literature searches via the `artl` MCP^25^, an MCP server that provides scientific literature discovery and identifier translation capabilities using Europe PMC^26^. It allows the curator agents to search for papers by keyword, retrieve full metadata and content from identifiers, and convert papers to structured markdown. The curator agent then synthesises the retrieved literature into supporting citations and usage notes such as domain-specific usage patterns, and recommends the most appropriate ontology placement. It then returns a structured report including confidence levels, suggested definitions, synonyms, and candidate parent terms to the co-ordinating agent, which is responsible for high-level design decisions, for example whether to create the term in EFO or to re-use it from another ontology (Figure 1; Supplemental S3). The ontologist agent specialises in writing scientifically correct definitions, using the appropriate annotation property for each synonym, building relationships between terms, and broader editorial actions such as deprecating terms or adjusting the ontology hierarchy.

**Fig. 1:**
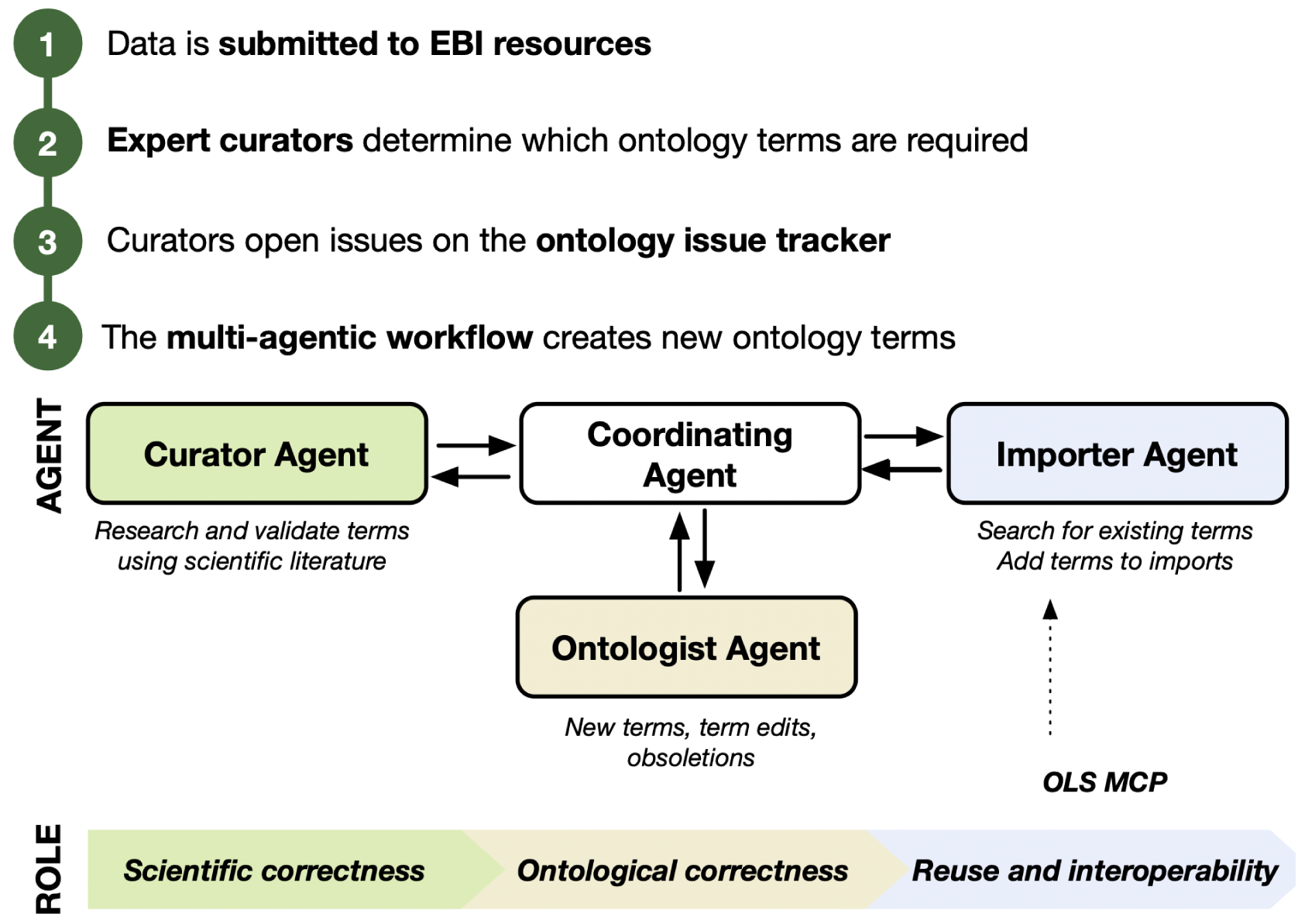
The multi-agent workflow deployed to enable automated updates to the Experimental Factor Ontology (EFO). The co-ordinating agent interprets issues opened by scientific curators, such as requests for new terms, term edits, and obsoletions which are handled by the ontologist agent. The curator agent is responsible for scientific correctness and researches and validates terms using the literature. The importer agent, responsible for reuse and interoperability, uses the OLS MCP to search for existing terms across different ontologies.

In the bronchiectasis example, curators needed to annotate data about specific endotypes of the disease. Endotypes classify diseases by distinct pathobiological mechanisms and can be used to stratify patients to deliver targeted therapies. For example, patients with an immune variant endotype may be treated with immune modulators such as GMCSF (granulocyte-macrophage colony-stimulating factor), while those with cilia variants as the predominant endotype may be targeted with motility enhancer drugs such as Sildenafil (Flume *et al.*, 2018). Curating this information, and subsequent emerging knowledge for example from future clinical trials, would require ontology terms for specific endotypes of bronchiectasis.

However, at the time of our bronchiectasis study, the only term available in EFO was the generic `MONDO:0004822 bronchiectasis`, so this information could not be grounded using ontology terms for specific endotypes. To address these missing terms, our agentic ontology workflow used the `artl` MCP to access the literature to identify a relevant publication, and proposed new ontology terms for each of the 8 endotypes (Supplemental S3). After validation by a human expert, these terms were included in the next EFO release to enable annotating data at the level of specific endotypes rather than only at the general level of the disease, including `EFO:0920040 ciliary dysfunction bronchiectasis` and `EFO:0920042 immune deficient-predominant bronchiectasis`.

To test the broader applicability of the MCP-grounded agents to general ontology editing, we used batch requests of 104 term imports and 5 new disease classes requested by users via the EFO issue tracker. Three expert ontology curators were assigned the same ticket. The same ticket was also separately assigned to our agentic workflow using Claude Sonnet 4.6 and the OLS MCP server. The agentic workflow completed the ticket overall faster than the human curators, at between 0:42 and 1:01 hours compared to between 1:18 and 2:55 hours. It should be noted that these durations are overall wall time and do not account for the variability of execution times of ontology development tools, and the human time included quality control which would still need to be performed on an AI curated output. We also compared six aspects of each pull request (PR) for inter-annotator agreement: labels, choice of parent classes, references, definition quality, clinical specificity, and conformance to EFO modelling conventions. All four PRs agreed on the preferred label for four out of five of the terms requested, and all four chose the same parental class for “refractory epilepsy”. Beyond these points of consensus, there were significant differences between each curator. For example, a term was requested for “bronze diabetes” which is an outdated name for the existing `EFO:1000642 hemochromatosis`, but only one human curator re-used the existing term with a new synonym rather than adding a new class, a mistake also made by the AI agent. The agentic workflow produced the most complete definitions across all five terms, consistently applying a genus-differentia definition structure, and included operational criteria for “refractory epilepsy” taken from the 2010 International League Against Epilepsy consensus. The agentic workflow also submitted 21 synonyms in total across all five terms, significantly higher than the average of two to three per human curator. Human curators made more minor errors, including typos (“dug-resistant” rather than “drug-resistant”), inconsistent use of imported ontologies (using a Human Phenotype Ontology term as a parent class for “bullous lung disease”); but humans contributed more thoroughly when it came to database cross-referencing, for example including cross references to terms in MeSH, MedGen, OMIT, and MPATH. Overall the variance between different human curators was similar to the variance between the human curators and the AI agent, and there remains a need for manual quality review in both human and AI curation.

Our agentic workflow has produced 97 terms reviewed by human curators and released in EFO version 3.90.0 (May 2026), and 27 terms currently undergoing review for the next EFO release as of June 2026. These terms include 13 new “blood metabolite level” and 14 “response to drug” terms, which will enable more granular data integration by avoiding the use of nonspecific terms in the GWAS Catalog. MCP-enabled agents for ontology curation are a key part of our AI semantics workflow, as they are able to both curate knowledge grounded in correct ontology terms via OLS, and create new ontology terms where they are not yet available. Human supervision and benchmarking are critical parts of our process to ensure agent behaviour is reviewed and benchmarked over time.

**Fig. 2:**
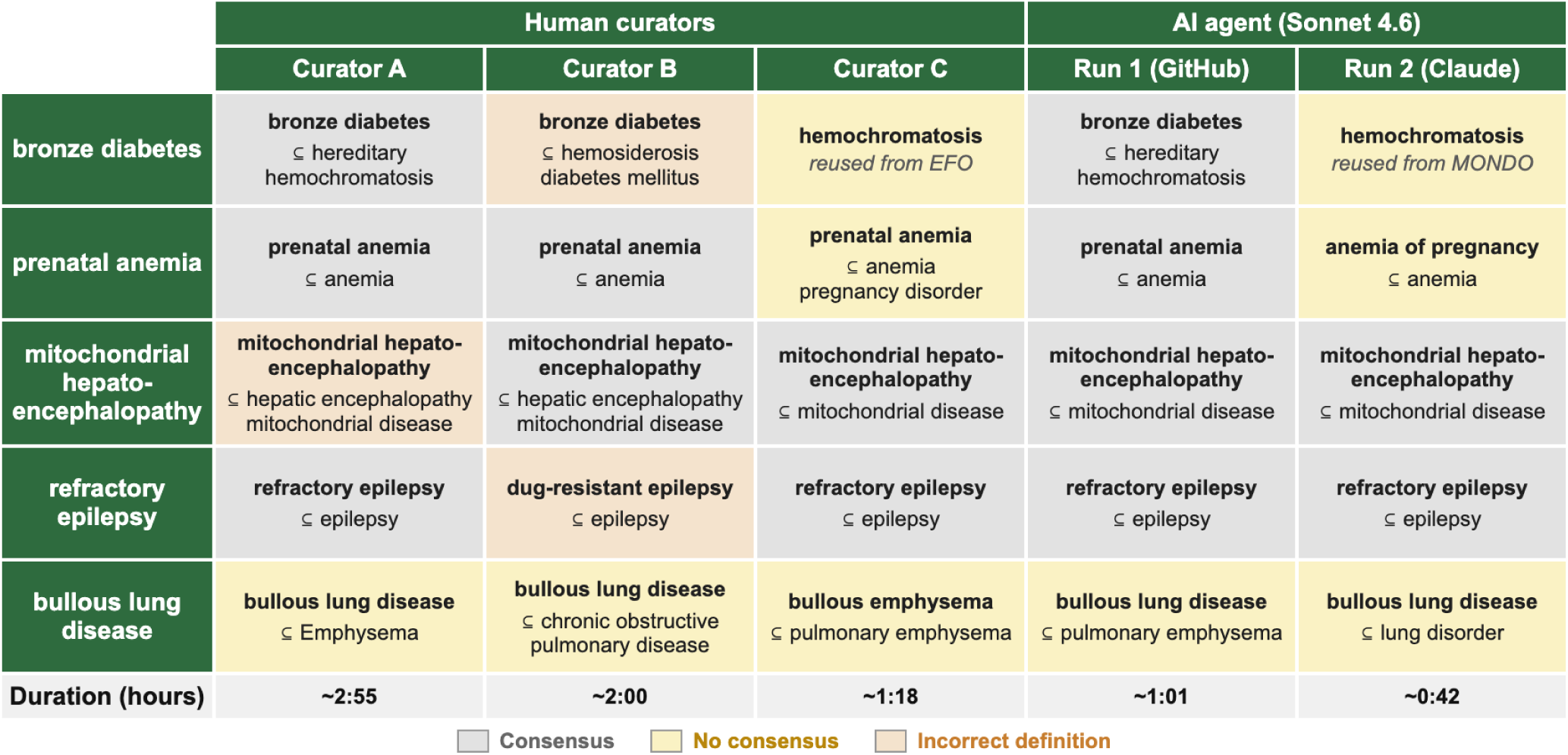
A comparison of parent terms, labels, and references for five terms added to the Experimental Factor Ontology (EFO) separately by three different human curators (A, B, C) and an AI agent (Sonnet 4.6, two separate runs via GitHub Copilot and Claude harnesses). Grey cells represent majority consensus; amber cells represent no consensus; red cells represent an incorrect definition.

### Scaling LLM embeddings to address semantic proliferation and promote semantic interoperability

Annotating data with ontology terms is part of the broader process of semantic alignment, which is necessary to make resources FAIR (Findable, Accessible, Interoperable, and Reusable)^27^. For databases to be interoperable, they must share semantics. Semantic alignment can include mapping from free text to standardised ontology identifiers, or mapping between semantically proliferated terms across different identifier spaces. For example, the text “increased liver size” maps to `HP:0002240 Hepatomegaly` used to annotate human phenotypes in the GWAS Catalog, which maps to `MP:0000599 enlarged liver` used to annotate mouse phenotypes in IMPC. These terms have the same meaning, but are in different ontologies with different labels and identifiers and are therefore semantically proliferated. Mappings can also capture non-exact biological similarity. For example, while `HP:0004380 Aortic Valve calcification` and `MP:0006131 Calcified pulmonary valve` are different phenotypes in different species, both are calcification of heart valves, and associating them enables linking associated genes with similar presentation across human and mouse studies ^28^.

Recently LLM embeddings have been applied to enable semantic alignment even with limited or missing formal classification, by interpreting the cosine distance between embedding vectors as semantic distance between concepts^22^. Unlike ontology-based methods for similarity such as Jaccard distance, embedding distance does not rely on similar ontology classification to generate a similarity score, enabling any string or term to be compared in the context of the embedding model without the need for prior expert classification. For example, the measurement term `EFO:0004314 forced expiratory volume` does not share any ancestors and has no cross references to the phenotype term `HP:0030877 Reduced FEV1/FVC ratio`, but the labels have a cosine similarity of 0.78 using the `text-embedding-3-small` model.

We have scaled this approach to create publicly accessible embeddings of all ontologies in OLS (280 ontologies and ∼11 million terms as of July 2026) with four separate embedding models: `llama-embed-nemotron-8b, harrier-oss-v1-27b`, `text-embedding-3-large`, and `text-embedding-3-small.` The `text-embedding-3-*` OpenAI models were chosen for their wide availability and relatively low cost through the OpenAI API. `llama-embed-nemotron-8b` and `harrier-oss-v1-27b` were chosen based on their availability on HuggingFace making embeddings reproducible on local hardware, and high performance on the Massive Text Embedding Benchmark (MTEB) Leaderboard^29^. Generalist models were chosen over specific biomedical models both due to the wide variety of ontology terms in OLS, which are used for a holistic range of biomedically relevant annotation and are therefore not limited to strictly biomedical domain terminology, and due to previously shown high performance of generalist embedding models for short-context clinical semantic search^30^.

Curators searching for ontology terms in OLS can select between the pre-existing lexical search, and semantic search using an embedding model, which enables matching search terms based on semantic rather than lexical similarity. In an example of semantic search improvement over lexical mappings, if a curator performs an OLS lexical search for “CABG” (an acronym for coronary artery bypass graft), the top result is an unrelated result `dates(feed) c_ABGqdt` due to the lexical substring match in the identifier string (“`c_ABGqdt”)`, as of May 2026. Using semantic search with `llama-embed-nemotron-8b` (the first model provided by OLS for public search queries) the top result is `SNOMED:232717009 Coronary artery bypass graft` based on a semantic similarity score of 81.3% (Fig. 3). Semantic search also bridges across languages even when ontologies have not been translated; for example, `EFO:0004292 vaccination` is returned for εμβολιασμός (Greek) or 예방접종 (Korean), or vacunació (Catalan) using `llama-embed-nemotron-8b` even though only the English label is defined in the ontology, complementing recent translation efforts in ontologies such as HPO^31^.

**Fig. 3:**
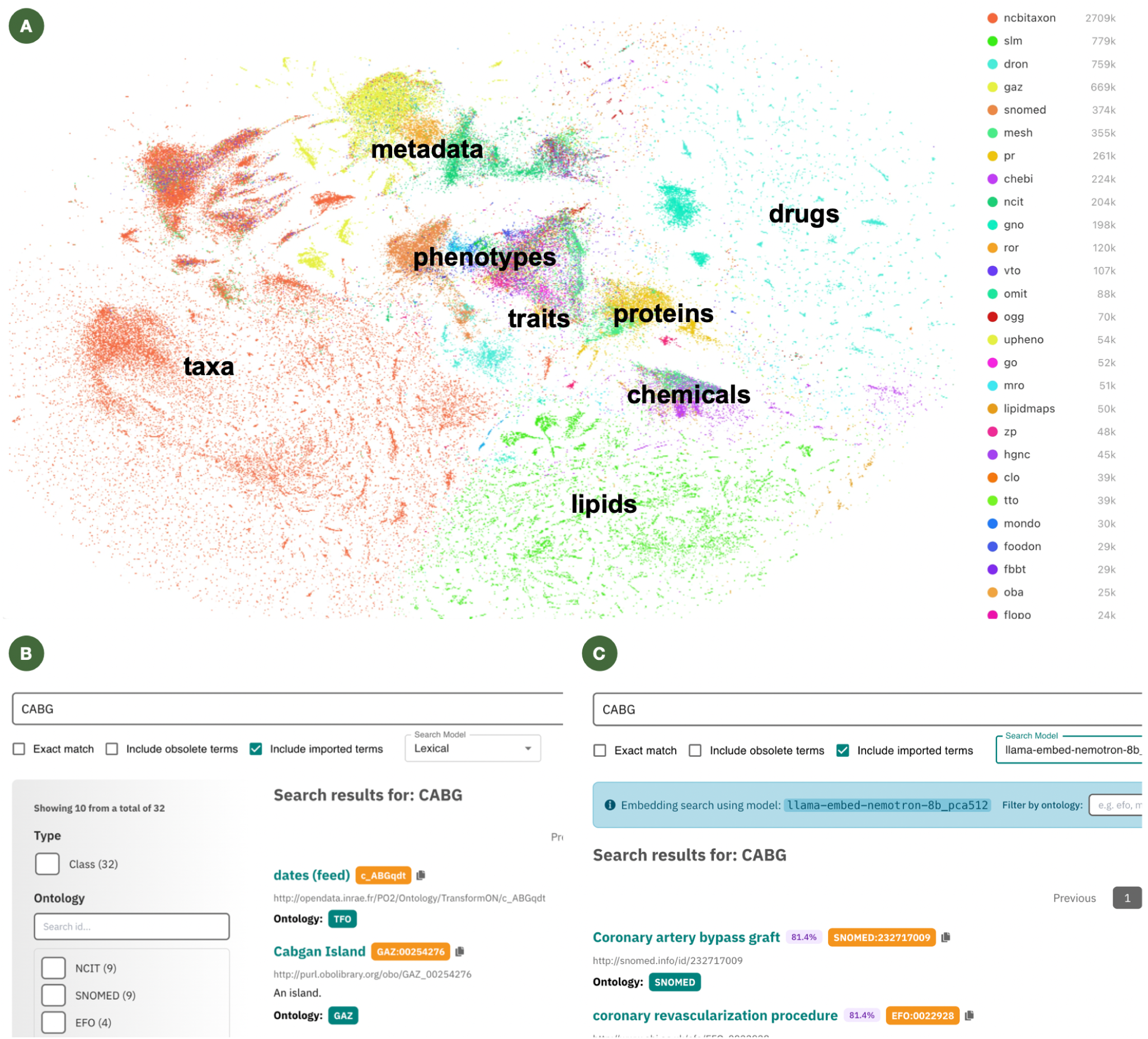
(a) Embeddings of all of the ontology terms in the Ontology Lookup Service (OLS) using `llama-embed-nemotron-8b` and visualised using UMAP. Each colour represents a different ontology, and each point represents a term. The distance between points visualises local structure of embedding space. Terms that are closer together are likely to be more semantically similar. This visualisation is embedded as an interactive widget on the OLS homepage. (b) Example of the pre-existing lexical search in the Ontology Lookup Service (OLS) for “CABG” (Coronary Artery Bypass Graft), returning an unrelated term “dates (feed)” as the top result due to the lexical substring match in the identifier string (“`c_ABGqdt”)`. (c) The new OLS semantic search for the same query “CABG” with the `llama-embed-nemotron-8b` model using embeddings. With semantic search, `SNOMED:232717009 Coronary artery bypass graft` is returned as the top result based on embedding distance.

In addition to providing semantic search using LLM embeddings, OLS now predicts similar terms based on embedding distance to map semantically similar by lexically different terms across ontologies. For example, the OLS term page for `EFO:0004309 platelet count` now displays links to proliferated (duplicated) measurement terms in different ontologies such as `OBI:2100012 platelet count assay` in the Ontology for Biomedical Investigations (OBI) used to annotate clinical EHR data^32^, even where these links are not defined in the ontology. The OLS API can be used to retrieve similar terms in bulk and at greater distances, including across species. For example, the `llm_similar` endpoint for `EFO:0004309 platelet count`, a term used to annotate human studies in the GWAS Catalog, includes the mouse phenotype `MP:0003179 thrombocytopenia` with a similarity score of 0.877 (https://www.ebi.ac.uk/ols4/api/v2/classes/http%253A%252F%252Fwww.ebi.ac.uk%252Fefo%252FEFO_0004309/llm_similar). New endpoints are also available to retrieve the cosine similarity between two classes (`llm_similarity`) and the underlying embedding vector for a class (`llm_embedding`). To enable large-scale mapping, all of the embeddings are also provided as Parquet files on the EBI FTP server, along with complete pairwise semantic similarity tables for selected disease and phenotype ontologies (currently EFO, HP, MP, OBA, MONDO, GO) to enable disease data integration.

We have evaluated the utility of embedding based similarity for cross-species mappings by comparing the results to existing manually curated HPO-MP mappings available in mapping-commons^33^, a consolidated repository of mappings derived from IMPC, Mouse Genome Informatics, and The Pistoia Alliance represented using the Simple Standard for Sharing Ontological Mappings (SSSOM)^34^. We evaluated the mean reciprocal rank (MRR) across the top 10 predicted HPO matches per MP term, restricting analysis to terms with `exact`, `equivalent` or `close` predicate relationships in the manually curated set (Fig 5a; Supplemental S4). MRR assesses how highly a model ranked the manually curated HPO-MP term pair among its candidate predictions. The highest MRR scores were achieved by `text-embedding-3-large` and `harrier-oss-v1-27b` followed by `text-embedding-3-small` and `llama-embed-nemotron-8b.` We also computed the recall at the top, top 5 and top 10 hits to evaluate each model’s capacity to identify the manually curated HPO-MP pair within its candidates. OLS embeddings using the `harrier-oss-v1-27b` model identified the correct mapping as the first hit in 81.4% of cases, `llama-embed-nemotron-8b` in 70.2%, `text-embedding-3-large` in 82.9%, and `text-embedding-3-small` in 78.8% of cases. (Fig. 4b). Notably, all models identified the correct HPO-MP mapping within their top 5 hits in >93% of cases, with the exception of `llama-embed-nemotron-8b` (87.9%). By the top 10 hits, all models identified the correct mapping in >91% of cases. Models with larger numbers of parameters performed better as expected; further investigation into inter-model differences would require analysis of training data which was not available for the four models tested.

**Fig. 4:**
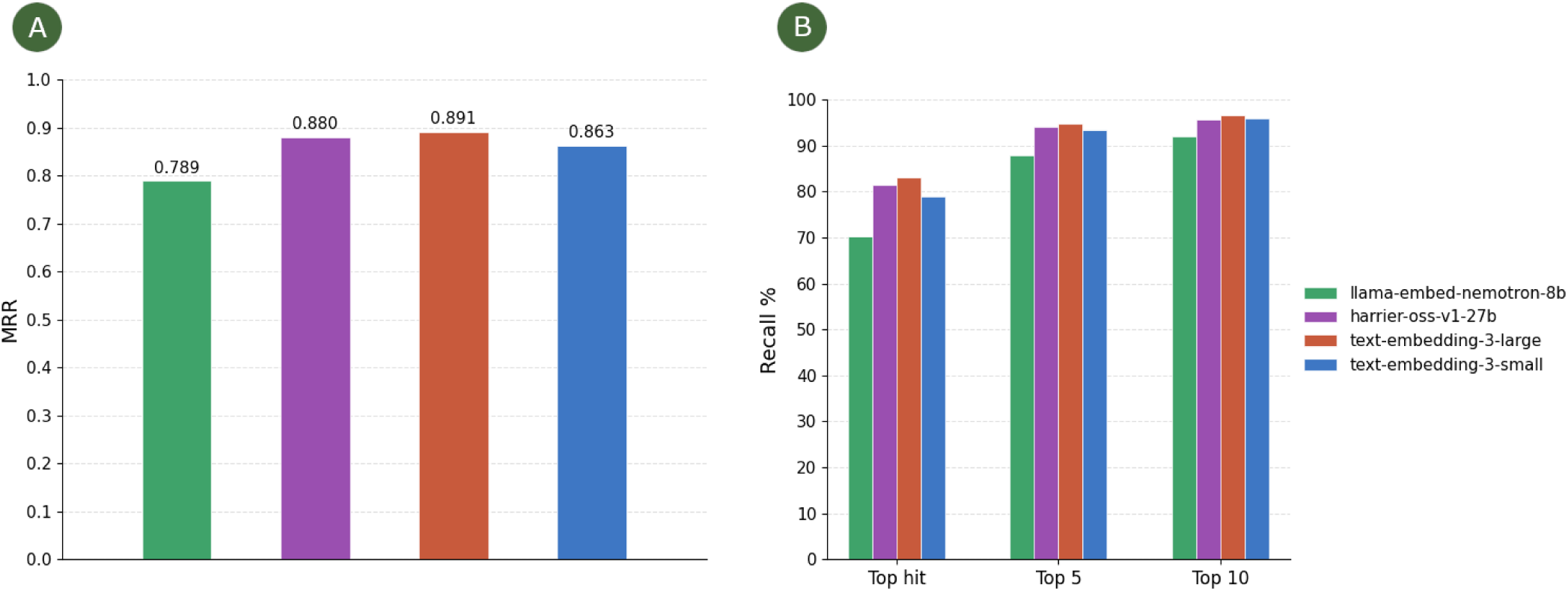
Evaluation of embedding-based mappings against expert-curated mappings between the Human Phenotype Ontology (HPO) and Mammalian Phenotype Ontology (MP). HPO-MP mapping embeddings produced with `llama-embed-nemotron-8b` (green), `harrier-oss-v1-27b` (purple), `text-embedding-3-large` (orange), `text-embedding-3-small` (blue) were evaluated against expert-curated reference HPO-MP mappings. Mappings were pooled by ontology predicates, including `skos:exactMatch, owl:equivalentClass,` and `skos:closeMatch` (N=2091). (a) Mean reciprocal rank (MRR) of the curated HPO-MP mapping among the top 10 candidate HPO terms predicted per MP term across all models. (b) Recall of expert-curated HPO-MP mappings found within the top hit, top 5, and top 10 of the embedding-based predictions.

**Fig. 5:**
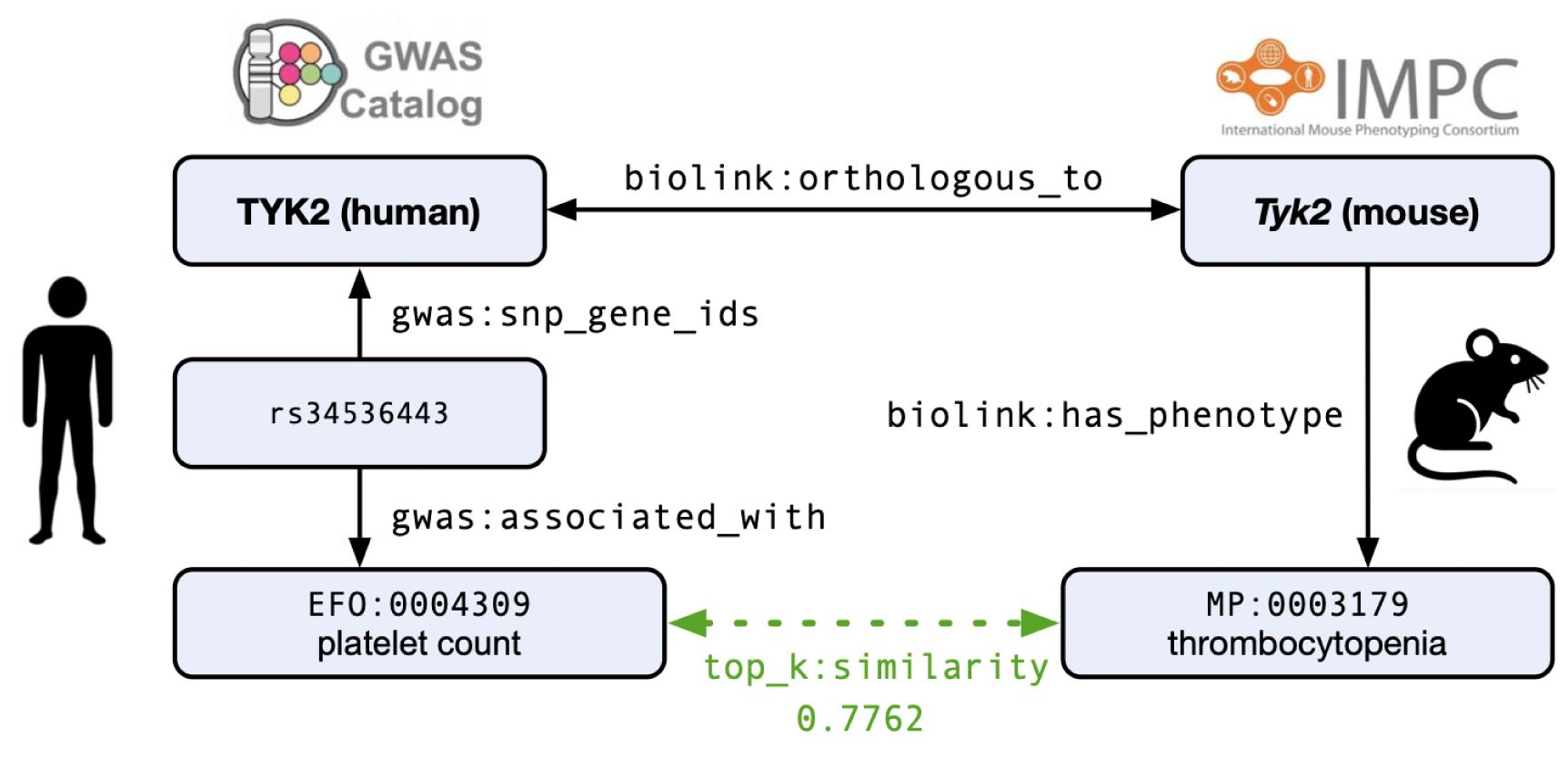
An example of orthologous genes which are also associated by similar phenotypes, by cosine distance between embeddings of their respective ontology terms in GrEBI. The human TYK2 gene is associated with `EFO:0004309 platelet count` in the GWAS Catalog, which has a similarity score of 0.7762 to `MP:0003179 thrombocytopenia` in IMPC using the OpenAI `text-embedding-3-small` model. The mouse Tyk2 gene, orthologous to human TYK2, is associated with thrombocytopenia in knockout mice.

OLS democratises the availability of embedding based methods for ontology mapping, by providing incrementally updated embeddings for all ontologies as part of the nightly OLS release cycle, along with standardised public APIs to access them without the need for an LLM provider or to a GPU to embed locally. As different models generate different results^35^ we have first benchmarked models, provided clarity on which model we have used, and documented the origin of embedding-derived mappings in our data and interfaces.

### Connecting data with LLM mediated knowledge graphs

The final component of our AI semantics workflow is GrEBI (Graphs@EBI), a knowledge graph and MCP server enabling LLM agents to perform queries across multiple biomedical databases including the GWAS Catalog, IMPC, Metabolights^36^, Reactome^37^, and MGnify^38^. Ontology terms used to annotate data are loaded with LLM embeddings from OLS, enabling queries across semantic and biological distance via embedding similarity of the ontology term labels and synonyms. For example, the human TYK2 gene is associated with `EFO:0004309 platelet count` in the GWAS Catalog, and the mouse ortholog Tyk2 is associated with `MP:0003179 thrombocytopenia` in IMPC. The GWAS Catalog and IMPC are different databases with different APIs and the ontology terms are not associated by cross-references, but are still connected in GrEBI when the embedding match is used to connect across the measurement and phenotype (Fig. 5), revealing shared phenotypes associated with orthologous genes.

The three components of our novel workflow (agentic curation with the OLS MCP, OLS LLM embeddings, and GrEBI) come together when a newly created ontology term is used as the entry point to integrate knowledge across resources (Fig. 6). As an end-to-end example, the agentic curation workflow created `EFO:0920021 cholesteryl esters in VLDL measurement`, to provide GWAS Catalog with a specific metabolite measurement term rather than relying on a more generic blood metabolite annotation. The GWAS Catalog data loaded into GrEBI links this measurement term to the SNP rs7412, which is linked to APOE in study `GCST90501287`, “A genetic map of human metabolism across the allele frequency spectrum” (p = 7 x 10^-307), The OLS term embeddings, also loaded into GrEBI, place the new term `EFO:0920021` close to `MP:0001550 hypercholesterolemia` and `MP:0001552 hypertriglyceridemia`, which are used as IMPC phenotype annotations for the mouse APOE ortholog Apoe (`MGI:88057`).

**Fig. 6:**
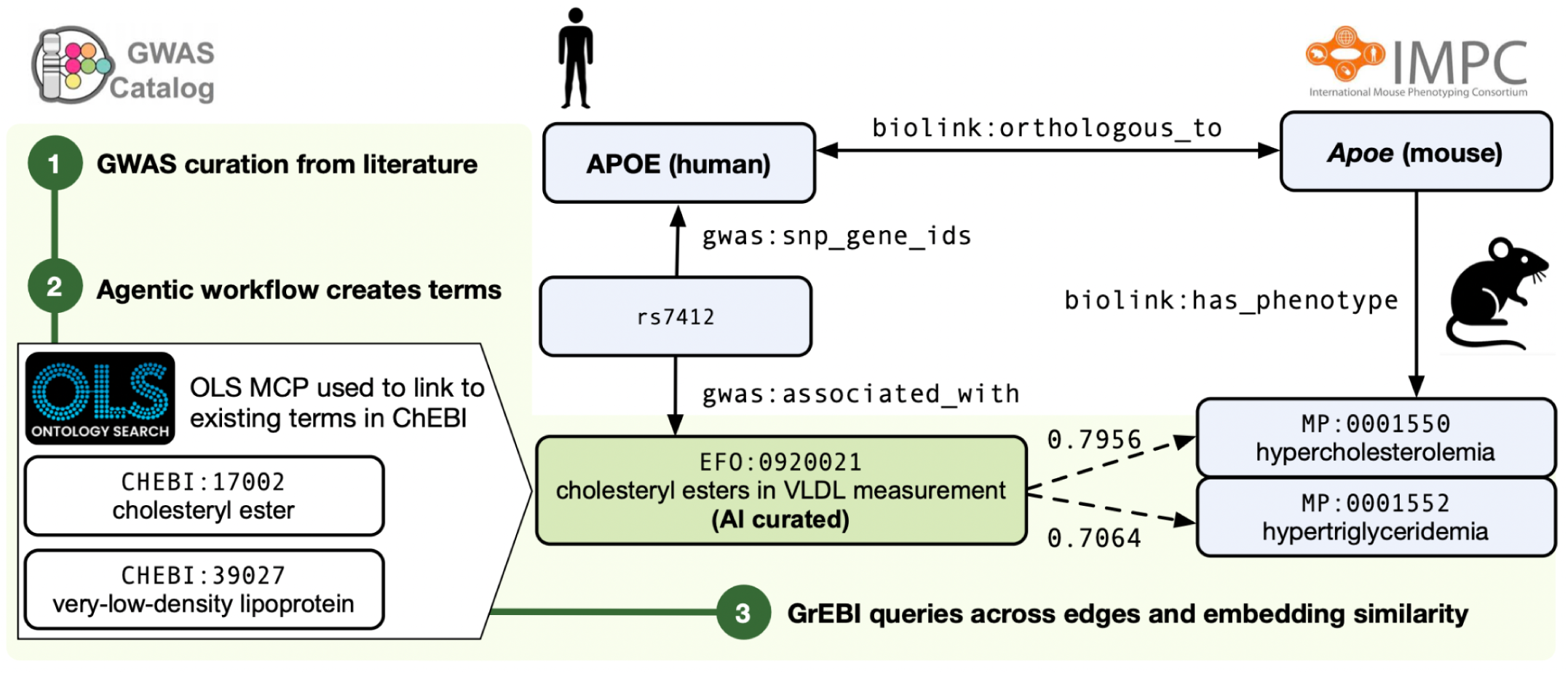
A complete iteration of our novel workflow for AI semantics. An expert curator in the GWAS Catalog requested a term “cholesteryl esters in VLDL measurement” which was created by our agentic workflow as `EFO:0920021`, using the OLS MCP to ground in existing ChEBI terms. The new term was automatically embedded by OLS, and has high semantic similarity to `MP:0001550 hypercholesterolemia` and `MP:0001552 hypertriglyceridemia` using the `text-embedding-3-small` model, which are phenotype terms used for mouse studies in IMPC which link to the mouse Apoe gene. Apoe is an ortholog of the human gene APOE, which contains a variant (rs7412) associated with the original GWAS trait. This example was discovered using the GrEBI KG, which incorporates all of the referenced ontologies, data, and embedding matches.

GrEBI is available to agents as a publicly accessible MCP server by defining Cypher graph query templates in YAML files, where each query has a defined set of parameters and result columns. For example, the `gwas_by_drugs_indicated_for_disease` query takes a `disease` identifier as an input parameter, and each result row has a `trait`, `drug`, `snp`, `gene`, `study`, and score. Providing controlled query templates ensures scalability of the underlying graph database by preventing unbounded queries from being executed. Each query template is also automatically used to define an MCP tool, such that adding a new query template is sufficient to provide a new capability to agents. This will enable GrEBI to scale to further knowledge bases across EMBL-EBI in the future.

GrEBI also generates REST API endpoints, and code examples in Python and R from the same templates. We have defined query templates across disease, phenotype, treatment, GWAS, metabolomics, and cross-species analysis use cases. We have also begun to apply the GrEBI MCP to complete the AI semantics workflow by directly informing the ontology editing agent with data content from biomedical databases. For example, to identify missing terms required for the GWAS Catalog we have created a `gwas_traits_reported_different_from_matched` query, which returns studies where the ontology term used for annotation does not have a matching label or synonym. When given access to the GrEBI MCP, the LLM agent was able to retrieve all traits mapped to `GO:0009410 response to xenobiotic stimulus` and any of its subclasses across ontologies. The coordinator agent was then able to create new terms following the same pattern as existing EFO terms related to drug responses. In future, this could enable a fully automated, closed loop curation workflow, in which the LLM agent creates new ontology terms as needed based on database content.

## Discussion

Our novel workflow demonstrates how LLMs and ontologies can be used synergistically to enable researchers to explore hypotheses across heterogeneous biomedical databases. We have demonstrated that limitations with existing ontology approaches - nonspecific and semantically proliferated ontology terms - can be addressed with LLMs, and conversely the major limitation of LLMs for ontology based integration (hallucinated or outdated terms) can be mitigated using grounding MCPs. The practical benefit of AI-assisted curation is the conservation and use of experts’ time. Human ontology curators are currently required for both routine tasks, such as extracting synonyms, generating consistent definitions, and importing terms from external ontologies; and decisions that require domain and ontology design expertise. We have shown that AI agents outperform human curators on routine tasks, suggesting our workflow can absorb the high-volume, structured components of curation as biological data grows in scale and complexity, enabling human experts to focus on more complex decisions that require biological and ontological domain knowledge.

Our approach assumes ontologies are trusted sources of ground truth for curation. This assumption is challenged when LLM agents participate in ontology development. While we have provided the agents with MCPs for grounding in correct identifiers and the scientific literature, an AI agent is still ultimately responsible for interpreting this information, and grounding does not equate to biological correctness; for example, `MONDO:0021163 kidney neoplasm` is a valid term to ground “kidney tumor”, but is not the same as `MONDO:0002367 kidney cancer`. Although in our EFO deployment all LLM assisted terms are reviewed by a human-in-the-loop, LLM hallucinations may be plausible to human expert curators. There is therefore a risk of these errors propagating undetected into downstream annotation, search, or data integration. Our current mitigation is to instruct the agents to provide explicit provenance reports, documenting how each definition statement and synonym is supported by primary literature, including the source publications and the specific sentences from which the information was derived. However, the ability of human curators to manually review every report is limited, and using agents for review risks the reports themselves containing hallucinated content. For transparency all terms added or updated by LLM agents have an annotation property (term editor “AI agent”), which will enable the use of more advanced LLMs in the future to validate ontology terms generated by older models, and could also be used to mitigate the risk of future LLMs being trained on LLM outputs by restricting training data to fully human curated terms. We plan to improve the granularity of this annotation by requesting extensions to the Evidence & Conclusion Ontology (ECO)^43^ to include terms for specific models. We also plan to implement real-time quality control by using multiple, separate AI agents in parallel with the same prompt, then comparing outputs by generating an inter-agent agreement score and before submitting for human review, in order to address the variance observed across both AI and human curators.

Similarly, in our embedding approach the main limitation is that unlike a human curator, OLS does not make a judgement on which is the most appropriate mapping, which is not always the closest match by embedding distance; for example, the top result for “mucus hypersecretion” in the OLS semantic search using llama-embed-nemotron-8b is `ZP:0015773 mucus secreting cell increased amount, abnormal` (score = 90.7%) which is a zebrafish phenotype, while the most appropriate match for annotating a human disease symptom would be `EFO:0005673 chronic mucus hypersecretion` (score = 83.4%). In future this could be mitigated by adding further context to refine the query (e.g. species), or introducing a post-processing step using an LLM agent to evaluate the top embedding matches and select the most appropriate, as in Retrieval Augmented Generation (RAG). Another limitation is that our current embedding strategy is restricted to labels and synonyms; in future we intend to evaluate enabling existing manually asserted classification to improve embedding recall by adding additional relationships such as ontology name, logical axioms, partonomy relationships used in anatomy ontologies, and applying a weighted average of embedding vectors with parent embedding vectors to inject ontology hierarchy.

Our workflow has been evaluated in the context of connecting disease data across species, but we have generalised all of our components to be broadly applicable to new biomedical data integration scenarios. For example, the OLS MCP server is published in BioContextAI^44^ where it is interoperable with a growing collection of MCP servers for application in agentic biomedical systems such as the Open Targets Platform MCP. Our agentic system for ontology development deployed for EFO has been adapted to develop a new ontology for identifying human cohorts (COHO). Our OLS embedding pipeline, originally evaluated in the context of HPO-MP mappings, extends to all of the ontologies in OLS, and has been used to map environmental exposures from the literature to the Environment Exposure Ontology (ECTO) as part of the EMBL Human Ecosystems transversal theme. Future work will include deploying GrEBI in both the GWAS Catalog and IMPC to provide enhanced graph-based search functionality taking advantage of our LLM-enriched ontologies, including searching GWAS by pathway, and to enable users to navigate between mouse and human studies based on shared phenotypes by embedding distance. In the GWAS case, we also plan to experiment with using the GrEBI MCP to fully automate deployment of needed ontology terms to re-annotate studies based on analysis of data content.

## Methods

Agents were implemented using the GitHub Copilot coding agent and custom Copilot agents to modularize domain-specific tasks. Custom agents were defined as Markdown agent profile files with YAML frontmatter specifying a unique name, description, model, and available tools, stored under `.github/agents/` in the ontology repository. The default Copilot coding agent served as the coordinating agent, driven by high-level instructions in `.github/copilot-instructions.md,` and was responsible for delegating subtasks to custom agents.

The OLS embeddings were performed on the EMBL-EBI high performance computing (HPC) cluster using a Nextflow pipeline. Local embeddings with `nvidia/llama-embed-nemotron-8b`^39^, and `microsoft/harrier-oss-v1-27b` were performed with the Sentence Transformers^40^ Python library. `text-embedding-3-large` and `text-embedding-3-small` embeddings were dispatched to the OpenAI API. Semantic similarity between terms was computed by comparing their embedding vectors using cosine similarity at thresholds of 0.4 and 0.7. Labels and synonyms are embedded separately to capture lexical variation.

Embedding mappings from HP version 2026-02-16 and MP version 2026-04-22 were evaluated against a manually curated reference set comprising 2091 unique HPO-MP term-pairs from three semantic relationship categories: `skos:exactMatch`, `owl:equivalentClass`, and `skos:closeMatch.` This reference set was assembled by merging mapping-commons^2^ files derived from IMPC, Mouse Genome Informatics, and The Pistoia Alliance^41^ (downloaded 2026-02-24). For each model, embedding term-pairs were ranked by cosine similarity and evaluated against the curated set using mean reciprocal rank (MRR) and recall. MRR was selected to evaluate ranking performance among the top 10 candidates for each MP term. Recall was computed to measure the proportion of manual mappings recovered within the top 1, top 5, and top 10 hits per category.

GrEBI (Graphs@EBI) was implemented using a Nextflow pipeline, with Rust and Python programs for the extract, transform, load (ETL) process. The GWAS Catalog version was 2026-04-27, IMPC DR 23.0, EFO version 3.88.0, HPO version 2026-02-16, and MP version 2026-03-24. All upstream data sources were converted to JSON objects for ingest. Identifiers were harmonised using the Bioregistry^42^ using prefix matching to canonicalise on a common prefix. Cliques of exact matches were merged using SSSOM^34^ mapping tables, and field values which match identifiers of other objects were linked as navigable edges in the graph. The knowledge graph (24,556,762 nodes and 1,012,027,138 edges as of June 2026) was loaded into Neo4j 2026.05.0 Community Edition for querying.

## Data Availability

A snapshot of the LLM embeddings and GrEBI KG can be downloaded from the EBI FTP at https://ftp.ebi.ac.uk/pub/databases/spot/ai_semantics. The EFO ontology including the agentic workflow is available at https://github.com/EBISPOT/EFO. The OLS MCP server is accessible at https://www.ebi.ac.uk/ols4/mcp. The OLS LLM embeddings are accessible through the OLS user interface and API at https://www.ebi.ac.uk/ols4/. GrEBI is available via API and MCP server at https://www.ebi.ac.uk/spot/grebi. Source code is licensed using Apache 2.0 and is available at https://github.com/EBISPOT/ols4 and https://github.com/EBISPOT/GrEBI.

## Acknowledgements

JAM, APB, AI, ZMP, ES, AF, RW, FLG, LH, and HP were supported by EMBL core funds. JAM was supported by National Institutes of Health (NIH) [R24-OD011883, OT2OD033756]. JAM, AI, and NM were supported by NIH National Human Genome Research Institute Phenomics First Resource, NIH-NHGRI # 5RM1 HG010860, a Center of Excellence in Genomic Science. JAM, DP, RW, FLG, and DS were supported by National Institutes of Health [5UM1HG006370-14]. LH was supported by NHGRI Grant Number 1U24HG012542-01.

## Competing Interests

SK is an employee of AstraZeneca since 2022 and holds AstraZeneca stock options. BM is an employee of Pfizer UK since 2023 and holds Pfizer Inc stock options. YA is an employee of Syngenta.

## Notes

https://ftp.ebi.ac.uk/pub/databases/spot/ai_semantics

### Summary of Updates

Figures 3 and 4 merged into one figure with additional detail. Reformatted and competing interests statement added.

