## Supplementary material for "AI semantics for biomedical data integration": Supplemental S1.pdf

This is a copy of a shared ChatGPT conversation
**Report conversation**

What are the main phenotypes of Bronchiectasis? Use HPO terms

Here's a list of the **main phenotypes associated with Bronchiectasis**, expressed using **Human Phenotype Ontology (HPO)** terms:

### 🫁 Respiratory system

- **Bronchiectasis** — HP:0002110 *(defining feature)*
- **Chronic cough** — HP:0012735
- **Productive cough** — HP:0031245
- **Recurrent respiratory infections** — HP:0002205
- **Hemoptysis (coughing up blood)** — HP:0002105
- **Dyspnea (shortness of breath)** — HP:0002094
- **Wheezing** — HP:0030828
- **Respiratory distress** — HP:0002098
- **Pleuritic chest pain** — HP:0030820
- **Crackles (rales) on lung auscultation** — HP:0030822
- **Reduced pulmonary function** — HP:0006536
- **Obstructive lung disease** — HP:0006531

### 🌡️ Infectious / inflammatory features

- **Recurrent pneumonia** — HP:0006532
- **Chronic sinusitis** — HP:0011109
- **Nasal polyposis** — HP:0001743
- **Chronic rhinitis** — HP:0012384

### 💧 Sputum / secretion-related

- **Excessive sputum production** — HP:0031251
- **Purulent sputum** — HP:0031253
- **Foul-smelling sputum** — HP:0031252

### 💨 Systemic / secondary manifestations

- **Clubbing of fingers** — HP:0100759
- **Failure to thrive** — HP:0001508 *(in pediatric cases)*
- **Weight loss** — HP:0001824
- **Fatigue** — HP:0012378

### 🧬 Associated conditions (in syndromic or secondary bronchiectasis)

- **Kartagener syndrome** (includes **situs inversus totalis** — HP:0001696, **chronic sinusitis** — HP:0011109, **bronchiectasis** — HP:0002110)
- **Cystic fibrosis phenotype** — often includes **Pancreatic insufficiency** — HP:0001738 and **Meconium ileus** — HP:0001254

Would you like me to organize these into **major clinical clusters** (e.g., “infective,” “inflammatory,” “systemic”) or a **table linking each HPO term to common etiologies** (like cystic fibrosis, primary ciliary dyskinesia, immunodeficiency)?
