## Supplementary material for "AI semantics for biomedical data integration": Supplemental S4.ipynb

```
# %% [code]
import polars as pl
from oaklib import get_adapter
from pathlib import Path

# %% [markdown]
# HP-MP embedding benchmarking


# %% [markdown]
**Objective**: To estimate recall of HPO-MP embedding mappings against manually curated HPO-MP mappings. 
This currently includes, top hit, top 5 hits, top 10 hits. 

**Datasets**: 
- [Manual mappings](https://github.com/mapping-commons/mh_mapping_initiative)
- [HPO-MP llama-embed-nemotron-8b](https://ftp.ebi.ac.uk/pub/databases/spot/ols_embeddings/23-Feb-2025/embeddings/semsim/llama-embed-nemotron-8b_avg/hp_mp__llama-embed-nemotron-8b_avg__0.7.tsv.gz)
- [HPO-MP text-embedding-3-large](https://ftp.ebi.ac.uk/pub/databases/spot/monarch/text-embedding-3-large_avg/hp_mp__text-embedding-3-large_avg__0.4.tsv.gz)
- [HPO-MP text-embedding-3-small](https://ftp.ebi.ac.uk/pub/databases/spot/ols_embeddings/23-Feb-2025/embeddings/semsim/text-embedding-3-small_avg/hp_mp__text-embedding-3-small_avg__0.4.tsv.gz)
- [HPO-MP harrier-oss-v1-27b](https://ftp.ebi.ac.uk/pub/databases/spot/monarch/harrier-oss-v1-27b_avg/hp_mp__harrier-oss-v1-27b_avg__0.7.tsv.gz)


# %% [markdown]
## Paths

# %% [code]
# Define paths
# TODO: Adjust paths as needed

# Path to manually curated file repo and mappings dir
MANUAL_MAPPINGS = Path("mh_mapping_initiative","mappings")


# Embedding mapping paths
# ollama-nemotron
NEMOTRON_AVG = Path("hp_mp__llama-embed-nemotron-8b_avg__0.7.tsv.gz")


# Open AI 
# small-text
OPENAI_SMALL_AVG = Path("hp_mp__text-embedding-3-small_avg__0.4.tsv.gz")

# large-text
OPENAI_LARGE_AVG = Path("hp_mp__text-embedding-3-large_avg__0.4.tsv.gz") 

# microsoft harrier
HARRIER_AVG = Path("hp_mp__harrier-oss-v1-27b_avg__0.7.tsv.gz")


# Output 
# TODO: Make sure to create this directory
OUTPUT = Path("output")

# Path to write manual mapping file with unique hpo-mp term-pairs
MANUAL_MAPPING_UNIQUE_TERMS = Path(OUTPUT, "hpo-mp-manual-mapping-unique-terms.csv")


# %% [code]
# Use oaklib to homogenise the URIs
# Get adapter to retrieve MP ids from the embeddings files.
mp = get_adapter("sqlite:obo:mp")
hp = get_adapter("sqlite:obo:hp")

# %% [code]
# Read and vstack all relevant hpo-mp mappings available in the manual mappings directory

# IMPC - remove items where sssom:NoTermFound in object_id
impc_all = (
    pl.scan_csv(
        Path(MANUAL_MAPPINGS, "mp_hp_*_impc.sssom.tsv"),
        separator="\t",
        comment_prefix="#",
    )
    .collect()
    .select(
        [
            "subject_id",
            "subject_label",
            "predicate_id",
            "object_id",
            "object_label",
            "mapping_justification",
        ]
    )
    .filter(~pl.col("object_id").str.contains("sssom:NoTermFound"))
    .unique()
)

# pistoia - has extra col: confidence (we can remove) and lacks labels - add with oaklib
pistoia = pl.read_csv(
    Path(MANUAL_MAPPINGS, "mp_hp_pistoia.sssom.tsv"), separator="\t", comment_prefix="#"
)

# Temporary lf for label fetching because kernel crashes if all done in a single step
temp_pistoia_label = pl.LazyFrame(
    pistoia.select("object_id", "subject_id")
    .unique()
    .with_columns(
        subject_label=pl.col("subject_id").map_elements(
            lambda x: mp.label(x), return_dtype=pl.Utf8
        ),
        object_label=pl.col("object_id").map_elements(
            lambda x: hp.label(x), return_dtype=pl.Utf8
        ),
    )
)

# Join labels, deduplicate and select/reorder necessary columns
pistoia = (
    pistoia.join(
        temp_pistoia_label.select(["subject_id", "subject_label"]).collect(),
        on="subject_id",
        how="left",
    )
    .join(
        temp_pistoia_label.select(["object_id", "object_label"]).collect(),
        on="object_id",
        how="left",
    )
    .unique()
    .select(
        [
            "subject_id",
            "subject_label",
            "predicate_id",
            "object_id",
            "object_label",
            "mapping_justification",
        ],
    )
)


# mgi - has extra col confidence - reorder/select necessary columns
mgi = pl.read_csv(
    Path(MANUAL_MAPPINGS, "mp_hp_mgi_all.sssom.tsv"),
    separator="\t",
    comment_prefix="#",
).select(
    [
        "subject_id",
        "subject_label",
        "predicate_id",
        "object_id",
        "object_label",
        "mapping_justification",
    ],
)

# Merge into a single file and keep unique entries
# COMMENT: Column selection might neglect scores and comments and this might be relevant later on
pre_data = impc_all.vstack(mgi).vstack(pistoia).unique()

## Clean ids and labels
# Some of the files contain errors in labels and ids - this replacement dictionary should provide fixes for most. 
replacement_dict = {
    # There is a mistake in an MP:000095 -> it should be MP:0000955
    "MP:000095": "MP:0000955",
    # MP:0001782 is an alt id and URI is broken. Use MP:0000008 instead
    "MP:0001782": "MP:0000008",
    # MP:0007180 is most likely a typo for MP:0001780. Label suggests it is a related synonym.
    "MP:0007180": "MP:0001780",
    # MP:0001311 is an alt id and URI is broken: Use MP:0001852
    "MP:0001311": "MP:0001852",
    # MP:0000546 is an alt id and URI is broken: Use MP:0002109
    "MP:0000546" :"MP:0002109",
    # HP:0007222 is an alt id and URI is broken: Use HP:0007305
    "HP:0007222": "HP:0007305",

    # HP:0012274 (type of inheritance) should be HP:0012744 (femoral aplasia) according to the label. Mosty likely a typo 
    "HP:0012274": "HP:0012744"
}

# Replace items
data = pre_data.with_columns(
    pl.col('subject_id').replace(replacement_dict),
    pl.col('object_id').replace(replacement_dict)
)

# Replace curies as id's with URI's
data_uris = data.with_columns(
    object_id=pl.col('object_id').map_elements(lambda x: hp.curie_to_uri(x), return_dtype=pl.Utf8),
    subject_id=pl.col('subject_id').map_elements(lambda x: mp.curie_to_uri(x), return_dtype=pl.Utf8)
)

# Remove labels to keep unique mappings only. (Different labels might duplicate data unnecessarily)
data_uris_uniques = (
    data_uris
    .drop('mapping_justification', 'subject_label', 'object_label')
    .unique())

# Merge exact and equivalent for simplicity
exact_equivalent_df = (
    data_uris_uniques
    .filter(
        pl.col('predicate_id').str.contains_any(['owl:equivalentClass', 'skos:exactMatch'])
    )
    .with_columns(
        predicate_id=pl.lit("exact/equivalent")
    )
    .select('subject_id','predicate_id', 'object_id')
    .unique()
)

# Separate the exact and uniques, homogenise, re-join
data_uris_master = (
    data_uris_uniques
    .filter(~pl.col('predicate_id').str.contains_any(['owl:equivalentClass', 'skos:exactMatch']))
    .vstack(exact_equivalent_df)
)


# %% [code]
# Write manual mapping file
# data_uris_master.write_csv(MANUAL_MAPPING_UNIQUE_TERMS)

# %% [code]
# Reading embedding mappings

# TODO: Select the embedding mappings file to benchmark
MODEL = OPENAI_LARGE_AVG
# Set a threshold in case table is not filtered (For OpenAI use 0.4, for harrier and nemotron use 0.7)
threshold = 0.4

# NOTE: WARNING: FROM HERE THE PIPELINE ASSUMES: subject_id = MP and object_id = HPO - adjust `new_columns` to fit the right order.
mapping_embeddings = (
    pl.scan_csv(
        MODEL,
        separator="\t",
        new_columns=[
            "object_id",
            "object_label",
            "subject_id",
            "subject_label",
            "cosine_similarity",
        ],
    )
    .filter(pl.col("cosine_similarity") >= threshold)
    # Filter object_id and subject_id to include only HPO and MP
    .filter(
        pl.col("subject_id").str.contains_any(["MP_"]),
        pl.col("object_id").str.contains_any(["HP_",]),
    )
)

# %% [code]
# Group by subject_id, check scores across related object_ids keep the top hit, top 5, top 10
# Column to group by, default is subject_id (MP)
GROUP_BY_COL = 'subject_id'
GROUP_BY_ONT = "MP"

# Filter the embedding mappings to only MP terms that are in the mapping file
mapping_embeddings_clean = (
    mapping_embeddings
    .filter(
    pl.col('subject_id').is_in(data_uris_master['subject_id']))
)
# Sort by cosine similarity score, group_by COL and then slice by the top number needed.
# top 10
embeddings_top_10 = (
    mapping_embeddings_clean
    .sort('cosine_similarity', descending = True)
    .group_by(GROUP_BY_COL, maintain_order=True)
    .head(10)
    .collect(engine="streaming")
)

# top 5
embeddings_top_5 = (
    embeddings_top_10
    .sort('cosine_similarity', descending = True)
    .group_by(GROUP_BY_COL, maintain_order=True)
    .head(5)
)

# top
embeddings_top=(
    embeddings_top_5
    .sort('cosine_similarity', descending = True)
    .group_by(GROUP_BY_COL, maintain_order=True)
    .head(1)
)

# %% [code]
# Create sets for quicker lookups 
embeddings_top_set= set(
    zip(
        embeddings_top['subject_id'],
        embeddings_top['object_id']
    )
)

embeddings_top_5_set = set(
    zip(
        embeddings_top_5['subject_id'],
        embeddings_top_5['object_id']
    )
)

embeddings_top_10_set = set(
    zip(
        embeddings_top_10['subject_id'],
        embeddings_top_10['object_id']
    )
)

# %% [code]
# Helper function for lookups
def embedding_matches(subject_id: str, object_id: str, embedding_set: set) -> bool:
    return (subject_id, object_id) in embedding_set

# Test
embedding_matches(subject_id="http://purl.obolibrary.org/obo/MP_0003600",
                  object_id="http://purl.obolibrary.org/obo/HP_0000086", 
                  embedding_set=embeddings_top_set)

# %% [code]
# For all categories, calculate top, top 5, top 10 and write to a df

# Clean datafile name for logging:
model_clean = str("-".join(MODEL.parts[-2:]))
model_name_clean = model_clean.rstrip(".tsv.gz")

if threshold > 0.4:
    model_name_clean = model_name_clean.replace("__0.4",f"__{threshold}")

# Initialise a dataframe to store all model information.
model_stats = pl.DataFrame(
    schema={
        "evaluated_file": pl.String,
        "cs_threshold": pl.Float64,
        "predicate_id": pl.String,
        "top_hits_percentage": pl.Float64,
        "top_5_hits_percentage": pl.Float64,
        "top_10_hits_percentage": pl.Float64,
        "grouped_by_ontology": pl.String
    }    
)

# Categories to a list
predicate_ids = pl.Series(data_uris_master.select("predicate_id").unique()).to_list()

print(f"Grouped by: {GROUP_BY_ONT}")
print(f"File evaluated: {MODEL}")
print(f"Threshold: {threshold}")
print("---"*20)

# Start loop
for category in predicate_ids:
    print(category)

    # Create a final dataset to write at the end
    tmp_data = {}

    # Create a tmp df with the category we need, drop predicate and keep unique ids
    tmp_df = (
        data_uris_master.filter(pl.col("predicate_id") == category)
        .drop("predicate_id")
        .unique()
    )

    # Feature to skip matches that are in equivalent and exact
    if category not in ['exact/equivalent']:
           tmp_df = (
            tmp_df.join(
            exact_equivalent_df.select('subject_id','object_id'),
            on=['subject_id', 'object_id'],
            how='anti'
        )
        .unique()
    )
 
    # Compute the three values
    temp_top_matches = tmp_df.with_columns(
        pl.struct(["subject_id", "object_id"])
        .map_elements(
            lambda x: embedding_matches(
                x["subject_id"], x["object_id"], embeddings_top_set
            ),
            return_dtype=pl.Boolean,
        )
        .alias("match")
    )

    temp_top_5_matches = tmp_df.with_columns(
        pl.struct(["subject_id", "object_id"])
        .map_elements(
            lambda x: embedding_matches(
                x["subject_id"], x["object_id"], embeddings_top_5_set
            ),
            return_dtype=pl.Boolean,
        )
        .alias("match")
    )

    temp_top_10_matches = tmp_df.with_columns(
        pl.struct(["subject_id", "object_id"])
        .map_elements(
            lambda x: embedding_matches(
                x["subject_id"], x["object_id"], embeddings_top_10_set
            ),
            return_dtype=pl.Boolean,
        )
        .alias("match")
    )

    # Calculate scores:
    temp_matches_score = (
        round(temp_top_matches.filter(pl.col("match")).height / temp_top_matches.height * 100, 2)
    )
    temp_top_5_score = (
        round(temp_top_5_matches.filter(pl.col("match")).height
        / temp_top_5_matches.height
        * 100, 2)
    )
    temp_top_10_score = (
       round(temp_top_10_matches.filter(pl.col("match")).height
        / temp_top_10_matches.height
        * 100, 2)
    )

    # Save scores to a tmp dictionary
    tmp_data = {
        "evaluated_file": model_name_clean,
        "cs_threshold": threshold,
        "predicate_id": category,
        "top_hits_percentage": temp_matches_score,
        "top_5_hits_percentage": temp_top_5_score,
        "top_10_hits_percentage": temp_top_10_score,
        "grouped_by_ontology": GROUP_BY_ONT,
    }

    # Make results into a DF
    tmp_results_df = pl.DataFrame(tmp_data)

    # Stack results into the master df
    model_stats = model_stats.vstack(tmp_results_df)


    # Print category specific stats
    print(f"Predicate id: {category}")
    print(f"Percentage of top matches: {temp_matches_score}%")
    print(f"Percentage of top 5 matches: {temp_top_5_score}%")
    print(f"Percentage of top 10 matches: {temp_top_10_score}%")
    print(f"Total N per category: {tmp_df.height}")
    print("---"*20)

# Optional: check which terms are not in the embedding mapping files
# This operation is can be implemented lazily using an anti join
all_terms_unique = pl.LazyFrame(data_uris_master.drop("predicate_id").unique())

not_in_embeddings= (
      all_terms_unique.join(
            mapping_embeddings_clean.select("subject_id", "object_id"),
            on=["subject_id", "object_id"],
            how='anti'
      )
      .collect(engine="streaming")
)

# Collect general stats
total_mp_terms=all_terms_unique.unique(subset='subject_id').collect().height
total_hp_terms=all_terms_unique.unique(subset='object_id').collect().height
total_manual_mappings=all_terms_unique.collect().height
mappings_not_found=not_in_embeddings.unique().height  

# Print overall embedding mapping coverage 
print(f"Total number of MP terms: {total_mp_terms}")
print(f"Total number of HPO terms: {total_hp_terms}")
print(f"Total number of manual mappings: {total_manual_mappings}")
print(f"Number of manual MP-HPO mappings not found in the embedding mappings: {mappings_not_found}")


# Add to the df
model_stats = model_stats.with_columns(
      total_mp_terms=total_mp_terms,
      total_hp_terms=total_hp_terms,
      total_manual_mappings=total_manual_mappings,
      mappings_not_found=mappings_not_found    
)

# %% [code]
# Write file to output
# NOTE: This file will be needed to plot results in the Visualisation stage
OUTPUT_BENCHMARK = (Path(OUTPUT, f"{model_name_clean}-benchmark.csv"))
model_stats.write_csv(OUTPUT_BENCHMARK)

# %% [markdown]
## Visualisation

# %% [code]
# Combine dataframes as necessary:
# TODO: Adjust path and filename based on outputs from previous stage
df_1 = pl.read_csv(Path(OUTPUT, "text-embedding-3-small-hp_mp__text-embedding-3-small_avg__0.4-benchmark.csv"))
df_2 = pl.read_csv(Path(OUTPUT, "text-embedding-3-large-hp_mp__text-embedding-3-large_avg__0.4-benchmark.csv"))
df_3 = pl.read_csv(Path(OUTPUT, "nemotron-8b-hp_mp__llama-embed-nemotron-8b_avg__0.7-benchmark.csv"))
df_4 = pl.read_csv(Path(OUTPUT, "microsoft-hp_mp__harrier-oss-v1-27b_avg__0.7-benchmark.csv"))

# Vstack model files as needed
df_all_models = (
    df_1.
    vstack(df_2)
    .vstack(df_3)
    .vstack(df_4)
)

# Write data for all models if needed
# df_all_models.write_csv(Path(OUTPUT, "hpo_mp_all_models_benchmark.csv"))

# %% [code]
import matplotlib.pyplot as plt
import matplotlib.patches as mpatches

# ─── 1. Readable labels ───────────────────────────────────────────────────────
MODEL_LABELS = {

    # OPEN AI
    # Small
    "text-embedding-3-small-hp_mp__text-embedding-3-small_avg__0.4": "text-embedding-3-small",
    # Large
    "text-embedding-3-large-hp_mp__text-embedding-3-large_avg__0.4": "text-embedding-3-large",
    # OLLAMA-nemotron
    "nemotron-8b-hp_mp__llama-embed-nemotron-8b_avg__0.7": "llama-embed-nemotron-8b",
    # MICROSOFT-harrier
    "microsoft-hp_mp__harrier-oss-v1-27b_avg__0.7": "harrier-oss-v1-27b",

}

PREDICATE_LABELS = {
    "exact/equivalent": "Exact/Equivalent",
    "skos:closeMatch":     "Close",
    "skos:relatedMatch":   "Related",
    "skos:narrowMatch":    "Narrow",
    "skos:broadMatch":     "Broad",
}

plot_df = df_all_models.with_columns(
    pl.col("evaluated_file")
      .map_elements(lambda x: MODEL_LABELS.get(x, x), return_dtype=pl.Utf8)
      .alias("model"),
    pl.col("predicate_id")
      .map_elements(lambda x: PREDICATE_LABELS.get(x, x), return_dtype=pl.Utf8)
      .alias("predicate_label"),
)

# ─── 2. Config ────────────────────────────────────────────────────────────────
MODELS = [
    "llama-embed-nemotron-8b", 
    "harrier-oss-v1-27b",
    "text-embedding-3-large",
    "text-embedding-3-small", 

    ]

PREDICATES = ["Exact/Equivalent", "Close", "Related", "Narrow", "Broad"]

METRICS    = [
    ("top_hits_percentage",    "Top hit"),
    ("top_5_hits_percentage",  "Top 5"),
    ("top_10_hits_percentage", "Top 10"),
]

# Hue encodes model, shade encodes K (dark→light = R@1→R@5→R@10)
MODEL_COLORS = {
    "llama-embed-nemotron-8b": {
        "Top hit": "#046D38",
        "Top 5":   "#2CA05E",
        "Top 10":  "#7DD3A6",
    },
    "text-embedding-3-large": {
        "Top hit":  "#7F2704",
        "Top 5":  "#D94801",
        "Top 10": "#FD8D3C"
    },
    "text-embedding-3-small": {
        "Top hit": "#003D6D",
        "Top 5":   "#1E75B5",
        "Top 10":  "#6BAED6",
    },

    "harrier-oss-v1-27b": {
        "Top hit":  "#3D006E",
        "Top 5":    "#7B2FBE",
        "Top 10":   "#C084FC"
    },
}

BAR_W        = 0.10
MODEL_PAD    = 0.07
PRED_SPACING = 2.0

# ─── 3. Bar positions ─────────────────────────────────────────────────────────
cluster_w = len(METRICS) * BAR_W
group_w   = len(MODELS) * cluster_w + (len(MODELS) - 1) * MODEL_PAD

specs, tick_xs = [], []

for p_idx, pred in enumerate(PREDICATES):
    center      = p_idx * PRED_SPACING
    group_start = center - group_w / 2
    tick_xs.append(center)

    for m_idx, model in enumerate(MODELS):
        cluster_start = group_start + m_idx * (cluster_w + MODEL_PAD)
        row = plot_df.filter(
            (pl.col("predicate_label") == pred) & (pl.col("model") == model)
        )
        for k_idx, (col, label) in enumerate(METRICS):
            specs.append({
                "x":      cluster_start + k_idx * BAR_W + BAR_W / 2,
                "height": row.select(col).item(),
                "color":  MODEL_COLORS[model][label],
            })

# ─── 4. Draw ──────────────────────────────────────────────────────────────────
fig, ax = plt.subplots(figsize=(15, 6))

for s in specs:
    ax.bar(s["x"], s["height"], width=BAR_W, color=s["color"],
           edgecolor="white", linewidth=0.5)

ax.set_xticks(tick_xs)
ax.set_xticklabels(PREDICATES, fontsize=12)
ax.set_ylim(0, 105)
ax.set_ylabel("Recall (%)", fontsize=12)
ax.yaxis.grid(True, linestyle="--", alpha=0.4, zorder=0)
ax.set_axisbelow(True)
ax.spines[["top", "right"]].set_visible(False)
ax.set_xlabel("Predicate", fontsize=12)


# ─── 5. Table legend ──────────────────────────────────────────────────────────
metric_labels = [label for _, label in METRICS]

leg_ax = ax.inset_axes([0.85, 0.8, 0.15, 0.15])
leg_ax.set_xlim(0, len(metric_labels))
leg_ax.set_ylim(0, len(MODELS))
leg_ax.axis("off")

for j, mlabel in enumerate(metric_labels):
    leg_ax.text(j + 0.5, len(MODELS) + 0.1, mlabel,
                ha="center", va="bottom", fontsize=10, fontweight="bold")

for i, model in enumerate(MODELS):
    row_y = len(MODELS) - i - 1
    leg_ax.text(-0.05, row_y + 0.5, model,
                ha="right", va="center", fontsize=11, clip_on=False, fontweight="bold" )
    for j, mlabel in enumerate(metric_labels):
        leg_ax.add_patch(
            mpatches.Rectangle((j, row_y), 1, 1,
                               facecolor=MODEL_COLORS[model][mlabel],
                               edgecolor="white", linewidth=1)
        )


plt.tight_layout()
plt.show()

# %% [code]
# ─── Missed mappings per model ────────────────────────────────────────────────
missed_df = (
    plot_df
    .group_by("model")
    .agg(pl.col("mappings_not_found").first())
)

# Preserve MODELS order
missed_vals = {
    row["model"]: row["mappings_not_found"]
    for row in missed_df.iter_rows(named=True)
}

BAR_W_M = 0.5
xs      = list(range(len(MODELS)))

fig, ax = plt.subplots(figsize=(6, 5))

for i, model in enumerate(MODELS):
    ax.bar(
        xs[i], missed_vals[model],
        width=BAR_W_M,
        color=MODEL_COLORS[model]["Top hit"],
        edgecolor="white", linewidth=0.5,
    )
    ax.text(
        xs[i], missed_vals[model] + 0.3,
        str(missed_vals[model]),
        ha="center", va="bottom", fontsize=11, fontweight="bold",
    )

ax.set_xticks(xs)
ax.set_xticklabels(MODELS, fontsize=11, rotation=45, ha="center")
ax.set_ylabel("Mappings not found", fontsize=12)
ax.yaxis.grid(True, linestyle="--", alpha=0.4, zorder=0)
ax.set_axisbelow(True)
ax.spines[["top", "right"]].set_visible(False)
ax.set_xlabel("Model", fontsize=12)

plt.tight_layout()
plt.show()
```
