## Supplementary figures and images for "AI semantics for biomedical data integration"

### Supplemental S2.png

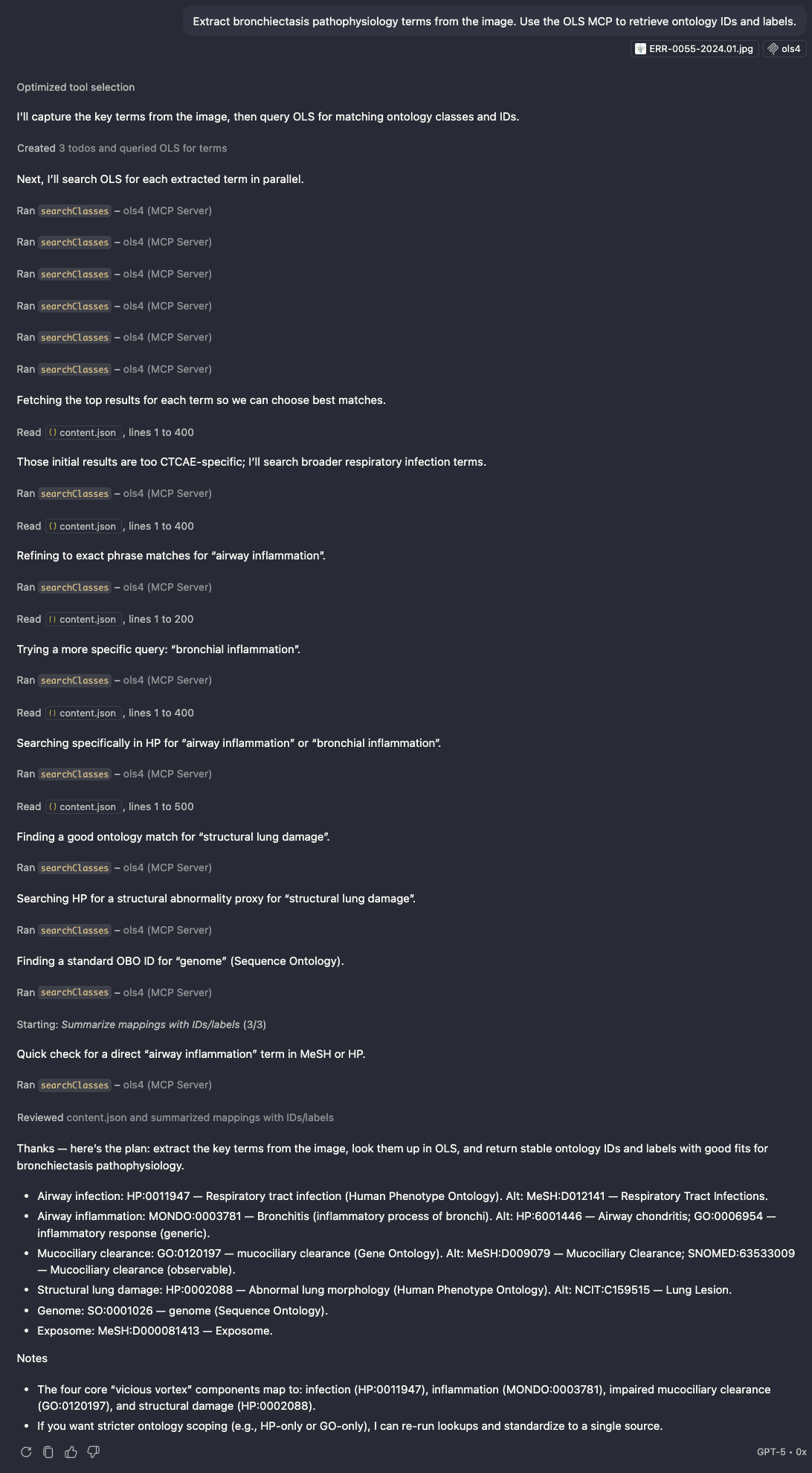
